# Structural and functional interrelationship of histone H2A and its variants H2A.Z and H2A.W in Arabidopsis

**DOI:** 10.1101/2024.12.03.626673

**Authors:** Youchao Wang, Jiabing Wu, Shuoming Yang, Xiang Li, Jiachen Wang, Qinghe Lv, Xiaoyu Zhu, Guoliang Lu, Jinru Zhang, Wen-Hui Shen, Bing Liu, Jinzhong Lin, Aiwu Dong

**Affiliations:** State Key Laboratory of Genetic Engineering, Institute of Plant Biology, Department of Biochemistry and Biophysics, School of Life Sciences, Fudan University, Shanghai 200438, P.R. China; State Key Laboratory of Genetic Engineering, School of Life Sciences, Zhongshan Hospital, Fudan University, Shanghai, P.R. China; State Key Laboratory of Genetic Engineering, Department of Biochemistry and Biophysics, School of Life Sciences, Fudan University, Shanghai 200438, P.R. China; Institut de Biologie Moléculaire des Plantes, CNRS, Université de Strasbourg, 12 rue du Général Zimmer, 67084 Strasbourg Cédex, France; Shanghai Institute of Infectious Disease and Biosecurity, Fudan University, Shanghai, P.R. China; Center for mRNA Translational Research, Fudan University, Shanghai, P.R. China

## Abstract

Multiple histone H2A variants are known in eukaryotes. However, the functional relationship between H2A and its variants in plants remains largely obscure. Using CRISPR/Cas9 editing, we generated a mutant lacking four H2A isoforms in Arabidopsis and analyzed the functional and structural relationship between H2A and its variants H2A.Z and H2A.W. RNA-sequencing and phenotype analyses revealed mild changes in gene transcription and plant development in the mutants lacking H2A, H2A.Z, or H2A.W compared with the wild-type plants. Chromatin immunoprecipitation sequencing analysis showed that H2A is able to substitute for both H2A.Z and H2A.W across the genome, including in euchromatin and heterochromatin regions. However, H2A.Z replaced both H2A and H2A.W primarily within the euchromatin regions. By using DNA and histones derived from Arabidopsis, we constructed nucleosomes containing H2A, H2A.Z, or H2A.W and resolved their cryogenic electron microscopy structures at near-atomic resolution. Collectively, the results reveal the structural similarity and functional redundancy of H2A and H2A variants in Arabidopsis.

## Introduction

In eukaryotes, the assembly of DNA into chromatin is crucial for genome stability and basic DNA-related biological processes, including DNA replication, transcription, and repair. The fundamental unit of chromatin, the nucleosome, consists of 146–147 bp DNA and a histone octamer, which contains a tetramer of histone H3–H4 and two dimers of H2A–H2B (Luger et al., 1997). Core histones are the most conserved proteins in eukaryotes and, in addition, eukaryotes have evolved non-allelic isoforms of core histones, known as histone variants (Malik and Henikoff, 2003; Talbert et al., 2012). The incorporation of histone variants within the nucleosome alters nucleosome stability, histone–DNA interaction, and the accessibility of chromatin-binding proteins, thereby influencing the DNA-based processes (Probst, 2022; Wu et al., 2024). In flowering plants, three H2A variants are known: H2A.Z, H2A.W, and H2A.X (Kawashima et al., 2015). H2A.Z and H2A.X exist conservatively in eukaryotes, whereas H2A.W is a plant-specific H2A variant (Kawashima et al., 2015).

H2A.Z is highly conserved among eukaryotes, exhibiting sequence conservation of approximately 70%–90% (Iouzalen et al., 1996). H2A.Z differs from H2A mainly in the amino acids within the L1 loop, the docking domain, and the acidic patch (Kawashima et al., 2015; Foroozani et al., 2022). H2A.Z is associated with diverse biological processes in plants, including regulation of flowering time (Wu et al., 2023), maintenance of genome stability (Rangasamy et al., 2003), DNA repair (Rosa et al., 2013), stress response and environmental adaption (Sura et al., 2017; Xue et al., 2021), and plant immunity (March-Diaz et al., 2008). H2A.Z is primarily enriched at the transcription start site and in the gene body in plants (Du et al., 2020; Wu et al., 2023). In Arabidopsis, three isoforms of H2A.Z are encoded by the genes *H2A.Z.8*, *H2A.Z.9*, and *H2A.Z.11* (Yi et al., 2006). The triple mutant lacking *H2A.Z.8*, *H2A.Z.9*, and *H2A.Z.11* is viable and shows multiple but mild changes in phenotypes, such as early flowering, smaller rosette leaves, stunted growth, weak gravitational response, and more lateral roots (Sun et al., 2022).

H2A.W is an H2A variant unique to plants with an extended C-terminal tail (Yelagandula et al., 2014; Kawashima et al., 2015). H2A.W is mainly enriched in the constitutive heterochromatin regions (Yelagandula et al., 2014; Lorkovic et al., 2017; Jamge et al., 2023). The chromatin remodeler DECREASE IN DNA METHYLATION 1 (DDM1) is essential for the deposition of H2A.W at potential mobile transposable elements (Osakabe et al., 2021). Moreover, synthetic H2A with the KSPK motif of H2A.W specifically influences heterochromatin composition and function in fission yeast, mimicking the role of H2A.W in plant heterochromatin (Lei et al., 2021). Recently, the cryogenic electron microscopy (cryo-EM) structures of DDM1 in complex with a nucleosome containing H2A.W have been reported (Osakabe et al., 2024; Zhang et al., 2024).

To further investigate the functional relationship between H2A and its variants in plants, we generated a quadruple mutant lacking four genes coding for Arabidopsis H2A and a triple mutant lacking three genes coding for H2A.W. The deletion of H2A or H2A.W resulted in weak phenotypes compared with the wild-type plants. ChIP-seq analysis demonstrated that H2A and H2A.Z are able to substitute for each other and replace H2A.W. Furthermore, cryo-EM analysis indicated that the structures of nucleosomes containing H2A, H2A.Z, or H2A.W are similar, providing a structural basis in support of the functional redundancy among H2A and its variants in plants.

## Results

### Phenotypic analysis of mutants lacking H2A, H2A.Z or H2A.W in Arabidopsis

In Arabidopsis, four isoforms of H2A are encoded by genes *H2A.1*, *H2A.2*, *H2A.10*, and *H2A.13*, three H2A.Z by *H2A.Z.8*, *H2A.Z.9*, and *H2A.Z.11*, and three H2A.W by *H2A.W.6*, *H2A.W.7*, and *H2A.W.12* (Wu et al., 2024) (Supplementary Fig. S1). To investigate the biological roles of H2A, H2A.Z, and H2A.W, we edited four genes of H2A and three genes of H2A.W using CRISPR/Cas9 in the Col-0 ecotype, respectively. A homozygous quadruple mutant, designated *h2a*, was created by introducing a 1-base pair (bp) deletion in *H2A.1*, a 2-bp deletion in *H2A.2*, and 1-bp insertion in both *H2A.10* and *H2A.13*. Additionally, a homozygous triple mutant lacking three isoforms of H2A.W, named as *h2a.w*, was also obtained, which had a 32-bp insertion, a 2-bp deletion, and a 1-bp insertion in *H2A.W.6* within two designed targets, a 1-bp deletion in *H2A.W.7*, and an 88-bp deletion in *H2A.W.12* (Fig. 1, A and B). All these targeted mutations resulted in frameshift and premature stop codons. The previously described *h2a.z* triple mutant was included in the analysis (Sun et al., 2022). Western blotting experiments confirmed the specificity of antibodies against H2A, H2A.Z or H2A.W (Supplementary Fig. S2). Using the above specific antibodies, we further validated the absence of H2A, H2A.Z and H2A.W respectively in the *h2a*, *h2a.z*, and *h2a.w* mutants (Fig. 1C).

**Figure 1.**
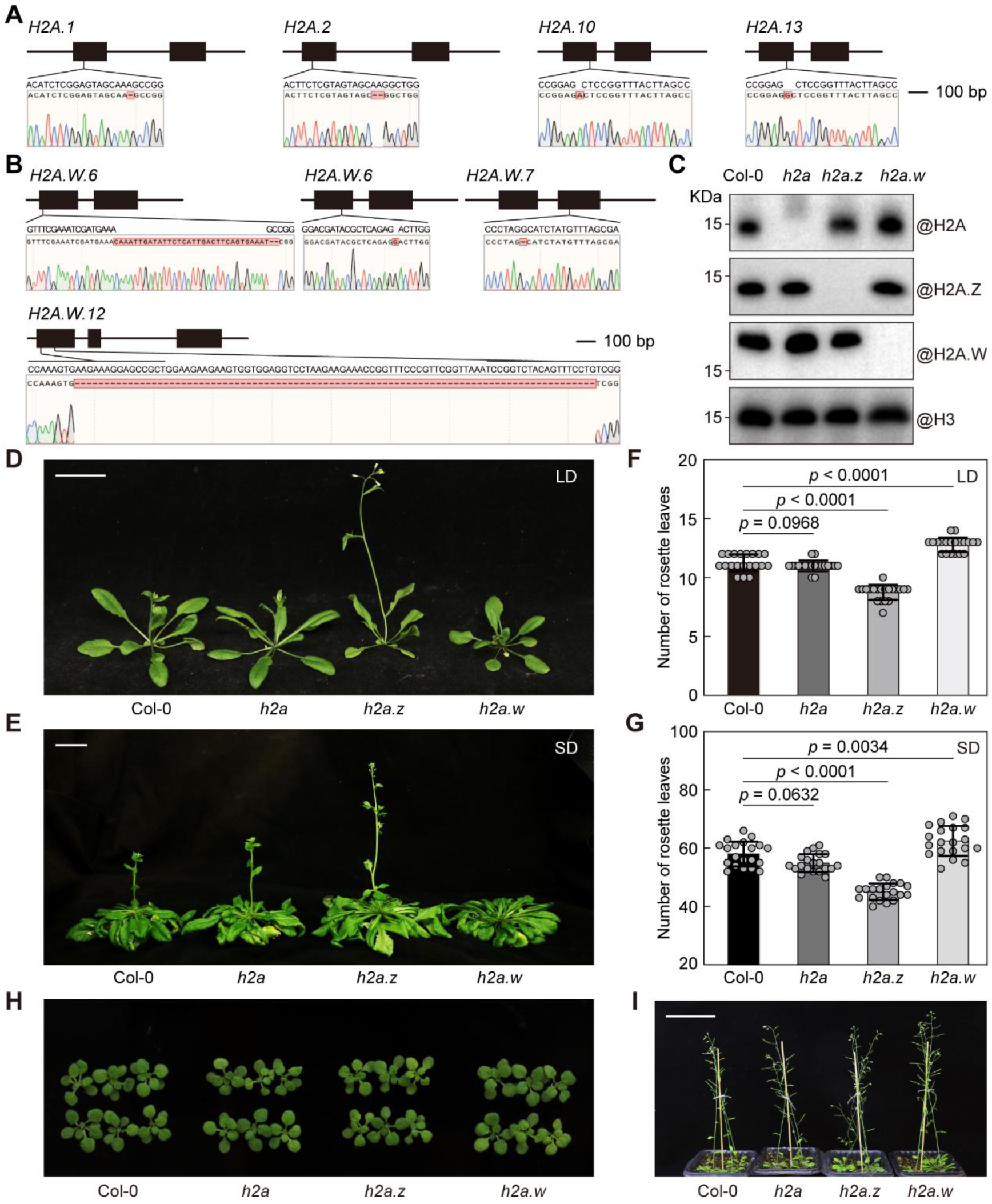
Loss of H2A, H2A.Z, or H2A.W does not cause severe developmental defects in Arabidopsis under normal conditions. **A, B)** Diagrams of the H2A or H2A.W coding genes showing base mutations in the *h2a* mutant (A) and *h2a.w* mutant (B). **C)** Western blotting showing the loss of H2A, H2A.Z or H2A.W in the indicated mutants, respectively. Nuclear extracts were analyzed using antibodies against H2A, H2A.Z, H2A.W, and H3. **D, E)** Representative images of *h2a*, *h2a.z*, *h2a.w*, and wild-type Col-0 seedlings grown under long-day (LD) (D) and short-day (SD) (E) photoperiod. Scale bar = 2 cm. **F, G)** Flowering time of *h2a*, *h2a.z*, *h2a.w*, and wild-type Col-0 plants grown under LD (F) and SD (G) conditions. Values shown are the mean ± standard deviation (SD) of 20 individual plants. Statistical significance was determined by a one-way ANOVA test. **H)** Representative images of *h2a*, *h2a.z*, *h2a.w*, and Col-0 seedlings grown under LD conditions for two weeks. Scale bar = 1 cm. **I)** Representative images of *h2a*, *h2a.z*, and *h2a.w*, and Col-0 seedlings grown under LD conditions for six weeks. Scale bar = 5 cm.

Phenotypic analysis revealed that the *h2a.z* and *h2a.w* mutants exhibited altered flowering times, with *h2a.z* showing early flowering and *h2a.w* displaying weakly delayed flowering (Fig.1, D–G). In contrast, the *h2a* mutant flowered similarly to the wild-type Col-0 (Fig.1, D–G). Further observations at both early and mature developmental stages showed that *h2a*, *h2a.z*, and *h2a.w* mutants were nearly indistinguishable from Col-0 in overall morphology (Fig. 1, H and I). The mild phenotype of the *h2a.w* mutant obtained in this study is similar to that previously reported (Bourguet et al., 2021). Therefore, the loss of *H2A*, *H2A.Z* or *H2A.W* does not cause severe developmental defects in Arabidopsis under normal growth conditions.

### Transcriptomic changes in *h2a*, *h2a.z*, and *h2a.w* mutants

To investigate how the loss of H2A or its variants affects gene transcription, RNA-sequencing (RNA-seq) was performed. Correlation analysis confirmed high reproducibility among the three biological replicates for each mutant (Supplementary Fig. S3). Genes encoding H2A and its variants, including *H2A.1*, *H2A.2*, *H2A.10*, *H2A.13*, *H2A.Z.8*, *H2A.Z.9*, *H2A.Z.11*, *H2A.W.6*, *H2A.W.7*, and *H2A.W.12*, showed the expected changes in *h2a*, *h2a.z*, and *h2a.w*, confirming the correction of these homozygous mutants (Supplementary Fig. S4). Differentially expressed genes (DEGs) were identified in each mutant: 513 genes in *h2a*, 743 genes in *h2a.z*, and 400 genes in *h2a.w* (Supplementary Fig. S5A, Supplementary Data Set 1), indicating that the deletion of H2A, H2A.Z, or H2A.W did not cause broad changes of gene transcription, which is consistent with the mild phenotypes observed in these mutants. Clustering analysis of DEGs showed that *h2a* and *h2a.z* exhibited similar transcriptional profiles, distinct from that of *h2a.w* (Supplementary Fig. S5B). To better understand the biological relevance of these transcriptional changes, we performed gene ontology (GO) enrichment analysis for the DEGs of each mutant. The GO analysis indicated that DEGs were mainly enriched in processes related to responses to environmental stimuli, particularly biotic and abiotic stresses (Supplementary Fig. S5, C and D), suggesting that H2A and its variants may contribute to plant adaptation to environmental changes.

### Replacement between H2A and its variants across the genome

To investigate genome-wide distribution changes following the loss of histone H2A or its variants, we performed chromatin immunoprecipitation sequencing (ChIP-seq) in *h2a*, *h2a.z*, and *h2a.w* mutants, as well as the wild-type Col-0 plants using antibodies respectively against H2A, H2A.Z, and H2A.W. Euchromatin and heterochromatin regions are marked by histone modifications H3K36me3 and H3K9me2, respectively (Liu et al., 2010; Roudier et al., 2011) (Fig. 2A). In Col-0, H2A was distributed across both euchromatin and heterochromatin regions. In contrast, H2A.Z was mainly enriched in euchromatin, whereas H2A.W was mainly located to pericentromeric heterochromatin regions (Fig. 2A).

**Figure 2.**
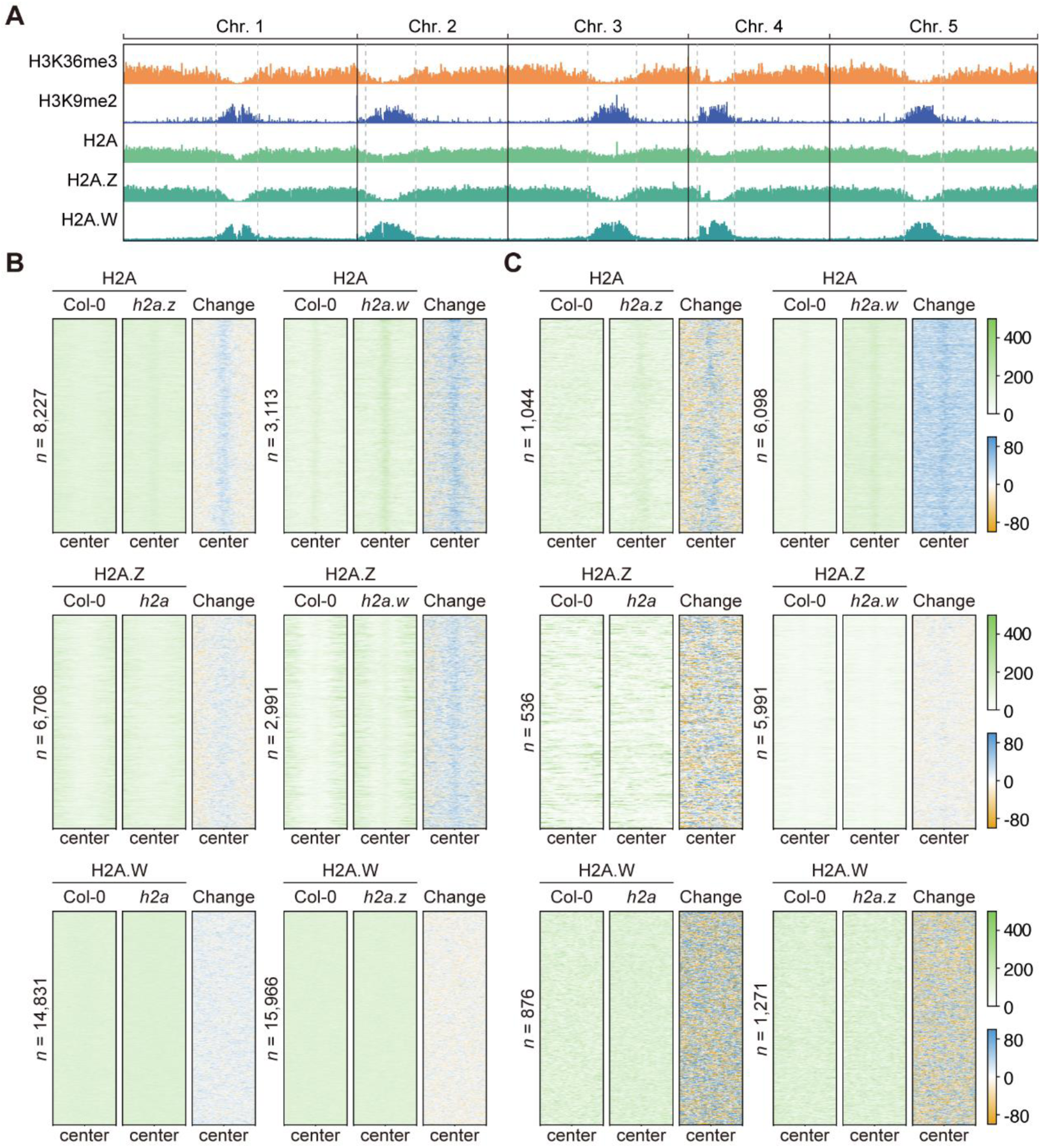
Genome-wide deposition and exchange among H2A, H2A.Z and H2A.W. **A)** Genome-wide distributions of H3K36me3, H3K9me2, H2A, H2A.Z, and H2A.W in the wild-type Col-0 Arabidopsis. **B, C)** Heatmaps showing the distributions and changes of H2A, H2A.Z, and H2A.W in Col-0 and the indicated mutants within the regions of chromosome arms (B) and pericentromeric heterochromatin (C) in the potential replacement peaks. All peaks are randomly sorted. Numbers of peaks are shown on the left.

To assess the distribution changes of H2A, H2A.Z, and H2A.W in the mutants, we only considered the newly appearing enrichment as a genuine increase in those histones compared to Col-0. In the *h2a.w* and *h2a.z* mutants, a significant increase in H2A was observed in both the euchromatic chromosome arms and pericentromeric heterochromatin regions (Fig. 2, B and C, Supplementary Data Set 2 and 3), suggesting that H2A can compensate for the absence of both H2A.Z and H2A.W. In contract, in the *h2a* and *h2a.w* mutants, H2A.Z enrichment was restricted to the chromosome arms and did not extend to the pericentromeric heterochromatin, suggesting that the replacement of H2A and H2A.W by H2A.Z is region-specific across the chromatin (Fig. 2, B and C). Notably, in the *h2a* and *h2a.z* mutants, no newly appearing enrichment of H2A.W was detected in either euchromatin or heterochromatin (Fig. 2, B and C). Taken together, our results demonstrated that Arabidopsis H2A is capable of replacing both H2A.Z and H2A.W across the genome, while H2A.Z is limited to replace H2A and H2A.W mainly within euchromatin regions.

### Screening plant native DNA for nucleosome reconstitution

To investigate the structural basis for the interchangeability of H2A and its variants, we reconstituted mononucleosomes using Arabidopsis-derived histones and DNA. Previously, nucleosomes were often assembled using the artificial Widom 601 DNA sequence, which does not accurately represent native DNA (Lowary and Widom, 1998). To better mimic physiological conditions, we screened multiple 147-bp DNA fragments from the Arabidopsis genome, each partially homologous (15 bp or 17 bp) to the Widom 601 sequence (Table 1). We evaluated these DNA fragments, designated as 17.1-147, 17.2-147, 15.1.1-147, 15.1.2-147, 15.2.1-147, and 15.2.2-147, for nucleosome assembly using Arabidopsis H2A, H2B, H3.1, and H4, expressed in *Escherichia coli* cells. Native-PAGE analysis showed that nucleosomes assembled with 15.2.2 fragment, hereafter referred to as 152 DNA, exhibited higher uniformity and stability comparable to those assembled with Widom 601 DNA (Fig. 3A). Therefore, the native 152 DNA fragment was selected for subsequent structural analyses.

**Figure 3.**
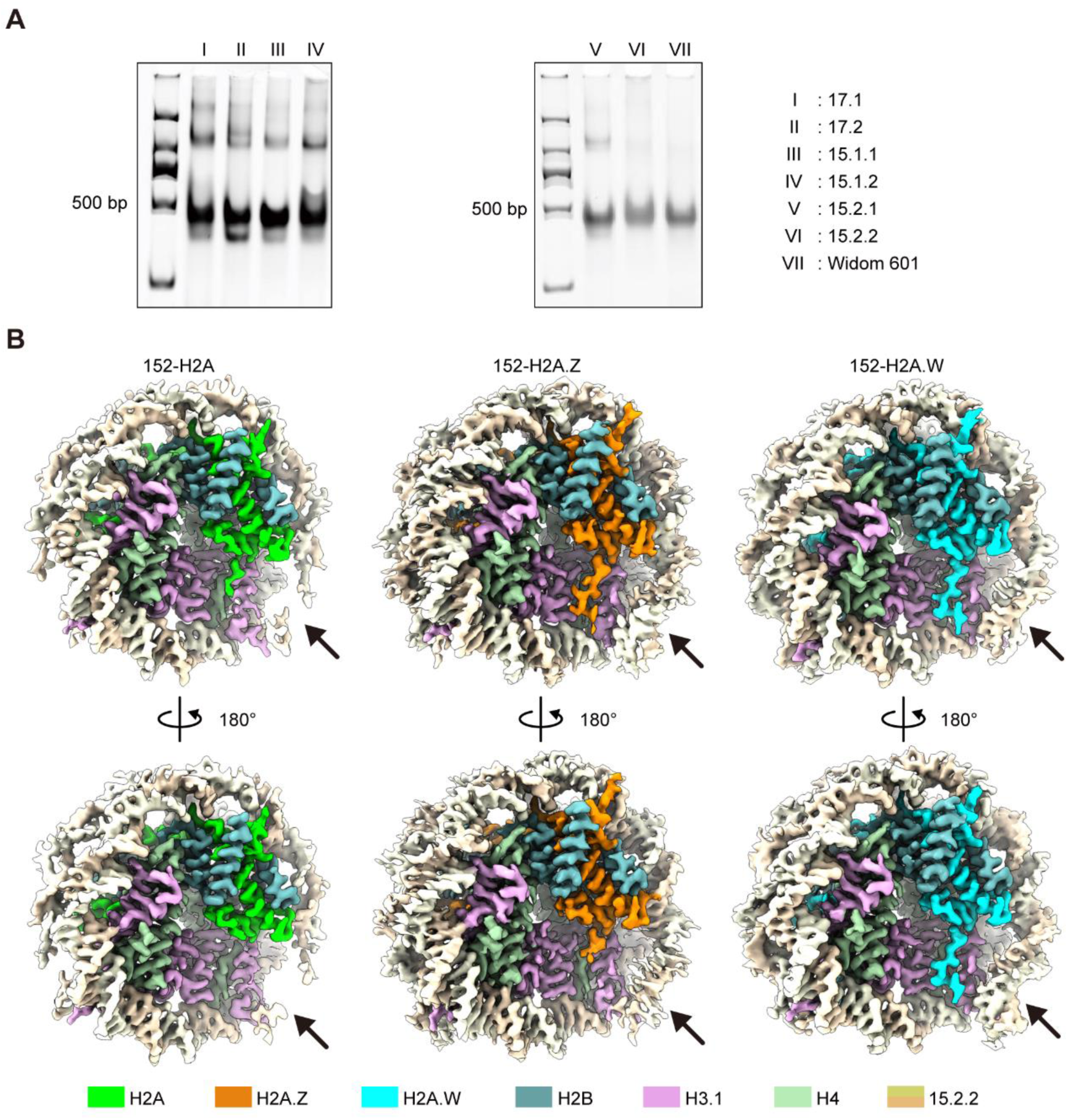
Cryo-EM structures of nucleosomes containing plant-derived DNA and histones. **A)** Arabidopsis DNA screening for mononucleosome reconstitution. Native-PAGE analysis of mononucleosomes assembled with seven different DNA fragments from Arabidopsis showing that nucleosomes assembled with the 15.2.2 (152) sequence display the highest uniformity among the DNA fragments tested, similar to the mononucleosomes containing Widom 601. **B)** Cryo-EM density maps imposed C1 symmetry of 152-H2A (left),152-H2A.Z (middle), and 152-H2A.W nucleosomes (right) at 3.34 Å, 2.93 Å, and 3.05 Å, respectively. The DNA entry/exit regions of the three nucleosomes are indicated by black arrows in each nucleosome.

**Table 1.**
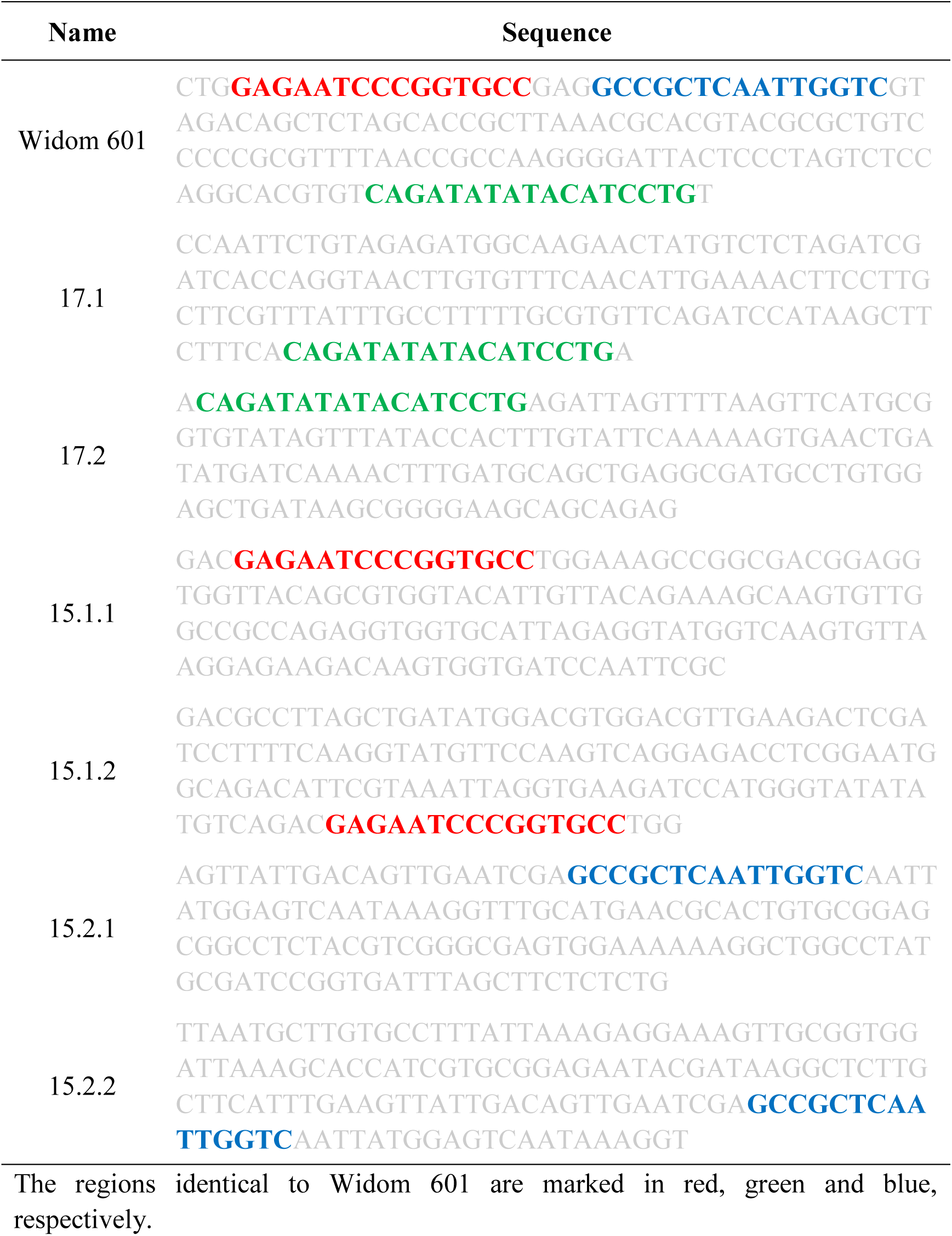
The sequence of DNA fragments derived from Arabidopsis used for mononucleosome reconstitution Name Sequence.

### Overall structures of mononucleosomes containing H2A, H2A.Z, or H2A.W

Using 300 keV cryogenic electron microscopy (cryo-EM), we resolved the structures of mononucleosomes containing Arabidopsis native 152 DNA and histones H2A, H2A.Z, or H2A.W, designated as 152-H2A, 152-H2A.Z, and 152-H2A.W, respectively (Table 2, Supplementary Figs. S6–S8). The cryo-EM maps based on C1 symmetry revealed that both ends of the DNA in H2A.W nucleosomes were well-resolved and rigid, whereas one DNA end in 152-H2A.Z nucleosomes was only partially resolved, and one end in 152-H2A nucleosomes was flexible and largely unresolved (Fig. 3B). The C1 maps achieved resolutions of 3.34 Å for 152-H2A, 2.93 Å for 152-H2A.Z, and 3.05 Å for 152-H2A (Fig. 3B).

**Table 2.** Cryo-EM data collection, refinement, and validation statistics for the nucleosomes in this study.

|  | 152-H2A<br>(PDB 9K40) | 152-H2A.Z<br>(PDB 9K3Z) | 152-H2A.W<br>(PDB 9K41) | 601-H2A<br>(PDB 9K42) | 601-H2A.Z<br>(PDB 9K43) |
| --- | --- | --- | --- | --- | --- |
| **Data collection and processing** |  |  |  |  |  |
| Microscope | Titan Krios | Titan Krios | Titan Krios | Titan Krios | Titan Krios |
| Camera | K3 Summit | K3 Summit | K3 Summit | K3 Summit | K3 Summit |
| Magnification | 64,000 | 81,000 | 81,000 | 81,000 | 81,000 |
| Voltage (kV) | 300 | 300 | 300 | 300 | 300 |
| Pixel size (Å) | 1.1 | 1.072 | 1.072 | 1.072 | 1.072 |
| Electron exposure (e<sup>-</sup>/Å<sup>2</sup>) | 50 | 50 | 50 | 50 | 50 |
| Exposure per frame (e<sup>-</sup>/Å<sup>2</sup>) | 1.25 | 1.25 | 1.25 | 1.25 | 1.25 |
| Number of frames collected | 40 | 40 | 40 | 40 | 40 |
| Defocus range (μm) | -0.9 to -1.8 | -0.9 to -1.8 | -0.9 to -1.8 | -0.9 to -1.8 | -0.9 to -1.8 |
| Symmetry imposed | C1, C2 | C1, C2 | C1, C2 | C1, C2 | C1, C2 |
| Micrographs (no.) | 1,112 | 1,116 | 1,521 | 1,609 | 2,695 |
| Initial particle images (no.) | 1,703,786 | 1,727,222 | 2,201,647 | 1,428,355 | 2,200,505 |
| Final particle images (no.) | 147,977 | 192,738 | 119,108 | 80,473 | 114,381 |
| Final reconstruction package | cryoSPARC | cryoSPARC | cryoSPARC | cryoSPARC | cryoSPARC |
| C1 Map resolution (Å) | 3.34 | 2.93 | 3.05 | 3.44 | 3.12 |
| C1 Map EMD ID | EMD-62035 | EMD-62033 | EMD-62037 | EMD-62039 | EMD-62041 |
| C2 Map resolution (Å) | 3.15 | 2.75 | 2.81 | 3.14 | 2.87 |
| C2 Map EMD ID | EMD-62036 | EMD-62034 | EMD-62038 | EMD-62040 | EMD-62042 |
| FSC threshold | 0.143 | 0.143 | 0.143 | 0.143 | 0.143 |
| **Refinement** |  |  |  |  |  |
| Refinement package | Phenix | Phenix | Phenix | Phenix | Phenix |
| Initial model used (PDB) | 1AOI | 1AOI | 1AOI | 1AOI | 1AOI |
| Map sharpening B factor (Å<sup>2</sup>) | -97.5 | -103.6 | -95.6 | -84.6 | -99.9 |
| **Model composition** |  |  |  |  |  |
| Non-hydrogen atoms | 11,824 | 11,814 | 11,828 | 11,824 | 11,814 |
| Protein residues | 746 | 744 | 746 | 746 | 744 |
| Nucleotide residues | 290 | 290 | 290 | 290 | 290 |
| Ligands | 0 | 0 | 0 | 0 | 0 |
| **R.m.s. deviations** |  |  |  |  |  |
| Bond lengths (Å) | 0.004 | 0.005 | 0.004 | 0.006 | 0.005 |
| Bond angles (°) | 0.764 | 0.756 | 0.736 | 0.958 | 0.755 |
| **Validation** |  |  |  |  |  |
| MolProbity score | 1.39 | 1.29 | 1.41 | 1.29 | 1.41 |
| Clash score | 6.99 | 5.44 | 6 | 5.36 | 7.42 |
| Rotamer outliers (%) | 0.65 | 0.64 | 1.29 | 0.49 | 0.8 |
| Cβ outliers (%) | 0 | 0 | 0 | 0 | 0 |
| CaBLAM outliers (%) | 0.84 | 0.98 | 0.56 | 0.84 | 0.56 |
| **Overall correlation coefficients** |  |  |  |  |  |
| CC (mask) | 0.84 | 0.78 | 0.8 | 0.82 | 0.81 |
| CC (box) | 0.82 | 0.69 | 0.75 | 0.82 | 0.76 |
| CC (peaks) | 0.78 | 0.63 | 0.67 | 0.76 | 0.7 |
| CC (volume) | 0.83 | 0.77 | 0.79 | 0.8 | 0.81 |
| **Ramachandran plot** |  |  |  |  |  |
| Favored (%) | 99.04 | 98.9 | 98.36 | 98.77 | 99.18 |
| Allowed (%) | 0.96 | 1.1 | 1.64 | 1.23 | 0.82 |
| Disallowed (%) | 0 | 0 | 0 | 0 | 0 |

To improve the resolution for histone octamer core within the nucleosomes, we applied C2 symmetry during reconstruction. This refinement improved the overall resolutions to 3.15 Å for 152-H2A, 2.75 Å for 152-H2A.Z, and 2.81 Å for 152-H2A.W (Supplementary Figs. S6D, S7D and S8D). The core region of the nucleosomes, particularly the histone octamer, showed higher resolution compared to the surrounding DNA regions (Supplementary Figs. S6F, S7F and S8F). To model these structures, we used the animal H2A nucleosome structure (PDB ID: 1AOI) as a reference, enabling precise atomic modeling of plant nucleosomes (Fig. 4A, Table 2, Supplementary Figs. S6G, S7G and S8G,).

**Figure 4.**
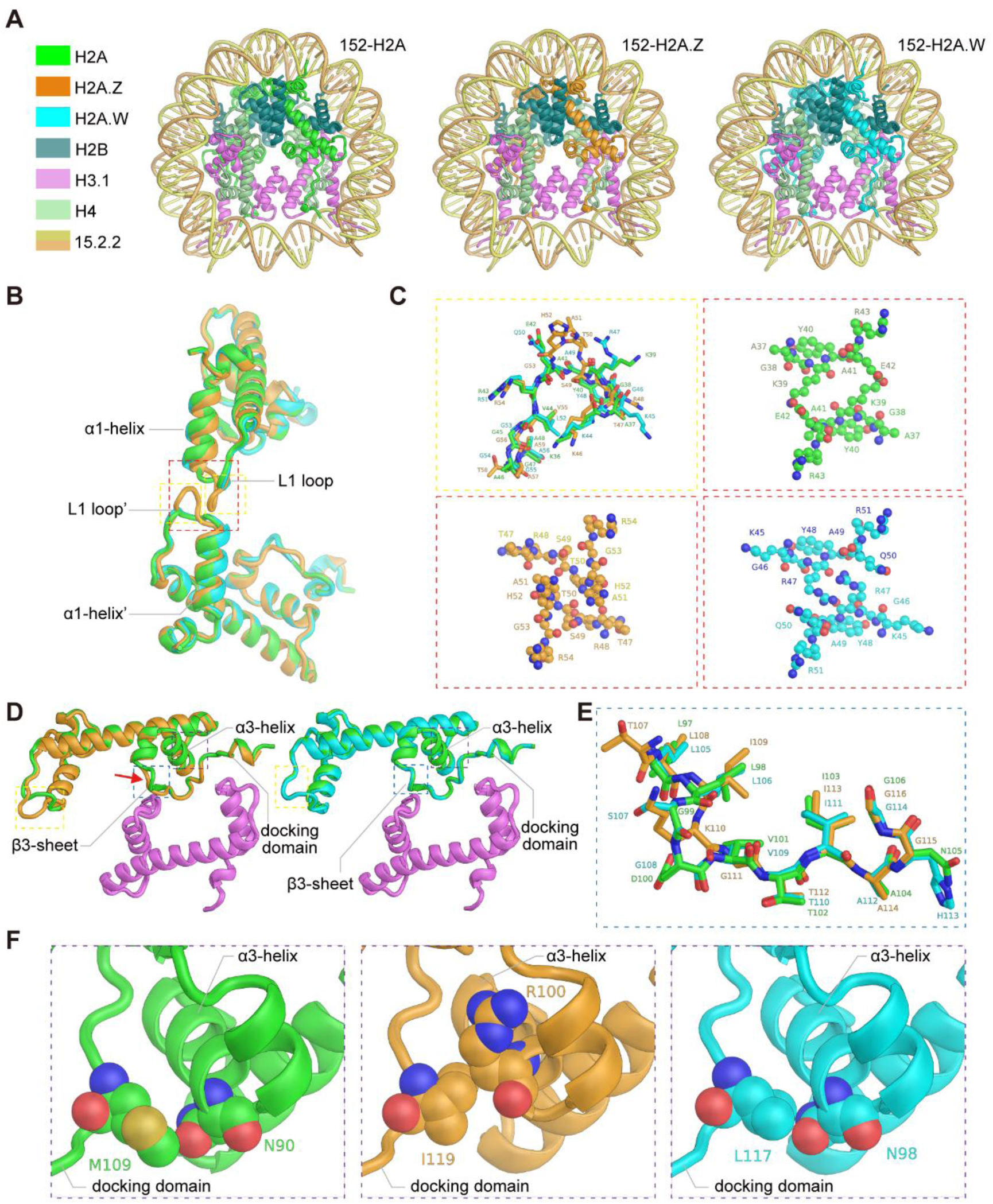
Structure comparison of nucleosomes containing Arabidopsis H2A, H2A.Z, or H2A.W and 152 DNA. **A)** Atomic models of 152-H2A (left),152-H2A.Z (middle), and 152-H2A.W nucleosomes (right) in disc view. **B)** Structure alignment of H2A, H2A.Z, and H2A.W interacting with their counterparts. **C)** L1 loop alignment of H2A, H2A.Z, and H2A.W shown as sticks (upper left). Two L1 loop interactions of H2A (upper right), H2A.Z (lower left), and H2A.W (lower right) with their counterparts shown as sphere. **D)** Structure alignment of H2A and H2A.Z complexed with H4 (left), H2A and H2A.W complexed with H4 (right), the differences of main chain between the two nucleosomes are indicated by red arrows. **E)** A part of the docking domain alignment for H2A, H2A.Z, and H2A.W shown as sticks. **F)** Close-up views of the purple dashed box from panel (D), showing internal interactions of H2A N90 and M109 (left); H2A.Z R100 and I119 (middle); and H2A.W N98 and L117 (right).

### Structural comparison of mononucleosomes containing H2A, H2A.Z, or H2A.W

The structures of the nucleosomes containing H2A, H2A.Z or H2A.W shared a largely conserved architecture, consisting of two H2A-H2B dimers and a (H3-H4)₂ tetramer that together formed the histone octamer, with approximately 146-bp DNA wrapped around it in a left-handed helix for about 1.65 turns. The charged surface representation of the histone octamer revealed variations in the acidic patches among the three nucleosomes, primarily due to differences in amino acid residues within H2A and its variants (Supplementary Fig. S1 and S9A). Structurally, H2A and H2A.W were nearly identical, while H2A.Z exhibits significant differences, particularly in its docking domain and L1 loop (Fig. 4, B–E, Supplementary Figs. S9B, S10, A and B). The L1 loop of H2A.Z extended further toward the other copy within the nucleosome compared to those of H2A and H2A.W (Fig. 4B). This extension increased the interacting interface and potentially strengthened the binding of the two H2A.Z subunits, leading to a tighter association between the α1-helix of two H2A.Z (Fig. 4, B and C, Supplementary Fig. S10A). In contrast, the binding of H2A and H2A.W with their respective counterparts was looser compared to that of H2A.Z (Fig. 4C, Supplementary Fig. S10A). It is noteworthy that the presence of the R47 residue in the H2A.W L1 loop increased the interacting interface between two H2A.W molecules, which may contribute to the increased rigidity of nucleosomes containing H2A.W (Fig. 4C, Supplementary Fig. S10A).

The docking domains of H2A, H2A.Z, and H2A.W all adopted a spoon-like shape, providing a large interface to interact with the (H3-H4)_2_ tetramer. The residues V101, T102, and I103 in H2A and V109, T110, and I111 in H2A.W formed a short β-sheet, referred to as β-3 domain. However, H2A.Z lacked the V residue within the region, which prevented the formation of the β-3 domain, leading to an altered orientation of the residue backbone in the docking domain of H2A.Z compared to H2A and H2A.W (Fig. 4, D and E, Supplementary Fig. S10B). Previous studies have showed that the rigidity of C-terminal docking domain of H2A contributes to the interaction between the H3 α-N helix and the DNA entry/exit regions (Armeev et al., 2021; Hirai et al., 2022; Osakabe et al., 2024). Our finding revealed that the interacting interface between L117 on the docking domain and N98 on the α3-helix of H2A.W was bigger than that of H2A (between M109 and N90), and that of H2A.Z (between I119 and R100) was intermediate (Fig. 4F, Supplementary Fig. S10C), which may explain the rigidity of the nucleosomes containing H2A, H2A.Z and H2A.W, respectively.

### Structural comparison of mononucleosomes containing plant native 152 DNA or artificial Widom 601 DNA

To compare the structures of nucleosomes reconstituted by plant native DNA with those by the artificial DNA, we assembled mononucleosomes using the 147-bp Widom 601 DNA and Arabidopsis histones, and further solved their structures by cryo-EM single particle assay (Table 2, Supplementary Figs. S11 and S12). Since no obvious differences were observed between 152-H2A and 152-H2A.W nucleosomes, we focused on assembling nucleosomes containing H2A and H2A.Z together with Widom 601 DNA, referred to as 601-H2A and 601-H2A.Z. The C1 map comparison revealed that the density map of 601-H2A was consistent with those of 601-H2A.Z and 152-H2A nucleosomes (Supplementary Fig. S13A).

The densities of 601-H2A and 601-H2A.Z were consistent with those of 152-assembled nucleosomes, exhibiting higher core resolution and lower resolution at the surrounding resolution (Supplementary Figs. S11F and S12F). We again used the animal nucleosome as a reference to build atomic models for the 601-assembled H2A and H2A.Z nucleosomes, achieving a good match of residues and nucleotides (Table 2, Supplementary Figs. S11G, S12G and S13B). The structure models of 601-H2A and 601-H2A.Z nucleosomes were identical to those assembled with 152 DNA (Supplementary Figs. S13, B–D). Furthermore, the DNA entry and exit ends of 601-H2A nucleosomes were consistently less rigid than those of 601-H2A.Z nucleosomes (Supplementary Fig. S13A).

Together, our structural analysis revealed that nucleosomes containing H2A or its variants display similar structures, supporting that H2A, H2A.Z, and H2A.W are capable to replace with one another.

## Discussion

In flowering plants, H2A, H2A.Z, and H2A.W show distinct genome-wide distributions. However, the mild phenotypes observed in the individual mutants in Arabidopsis suggest the functional redundancy of these histones. In this study, we observed that H2A can replace H2A.Z and H2A.W within the whole genome, while H2A.Z can substitute for H2A.W mainly within euchromatin but not pericentromeric heterochromatin regions. In addition, the cryo-EM structures of nucleosomes containing H2A, H2A.Z, or H2A.W are similar. Thus, our molecular and structural analyses support the functional redundancy of H2A, H2A.Z, and H2A.W in plants.

During the synthesis phase of the cell cycle, canonical histones, encoded by replication-dependent histone genes, are incorporated into chromatin (Marzluff et al., 2008). Given the crucial role of canonical histones in chromatin replication in eukaryotes, it is surprising that the mutant lacking four *H2A* genes displayed almost normal phenotypes compared with the wild-type Arabidopsis (Fig. 1, D–I). Compared with the roles of replication-dependent H3.1 and replication-independent H3.3, transcription linking with replication for H2A variants is less well understood in plants (Kawashima et al., 2015; Benoit et al., 2019). In yeast and animals, genes encoding canonical histones lack introns, which is a notable difference from those encoding histone variants and conductive to rapid, high-level expression of core histones (Marzluff et al., 2008). Interestingly, we observed that the genes encoding canonical H2A contain introns in flowering plants, whereas those encoding H2B, H3, and H4 lack introns (Supplementary Fig. S14), suggesting that plant H2A may function differently from those of yeast and animals.

The localization of H2A.Z and H2A.W is reported to be mutually exclusive: H2A.Z is predominantly associated with euchromatin, whereas H2A.W is enriched at heterochromatin (Wu et al., 2024). However, in the absence of H2A.W, H2A.Z can relocate to the position normally occupied by H2A.W within the chromosome arm regions in wild-type plants (Fig. 2, B and C). Instead, H2A can replace H2A.W in the pericentromeric heterochromatin (Fig. 2, B and C). Therefore, despite the similar structures of nucleosomes containing H2A, H2A.Z, and H2A.W, the replacement among H2A and its variants is position-specific *in planta*, which might be driven by the distinct positioning of histone remodeling factors or chaperones in plants (Du et al., 2020; Wu et al., 2023; Wu et al., 2024). The chromatin remodeler DDM1 is reported to interact with both H2A.Z and H2A.W through conserved binding sites, preventing H2A.W from being replaced by H2A.Z at the transposable elements (Jamge et al., 2023), which may explain why H2A.Z can replace H2A.W at the chromosome arms but not in the pericentromeric regions where DDM1 is located (Fig. 2, B and C).

The overall structure of mononucleosomes constructed by animal and plant H2A or its variants is similar (Luger et al., 1997; Lewis et al., 2021; Osakabe et al., 2024; Zhang et al., 2024), and subtle differences are observed in the L1 loop and docking domain within the 152-H2A.Z nucleosome compared with the 152-H2A and 152-H2A.W nucleosomes (Fig. 4, Supplementary Fig. S9). This is consistent with the differences between nucleosomes containing animal H2A and H2A.Z (Luger et al., 1997; Suto et al., 2000). The rigidity of the DNA end in plant nucleosomes containing H2A.Z is intermediate between that of the nucleosomes containing H2A or H2A.W (Fig. 3B). By contrast, the DNA entry and exit sites in animal nucleosomes containing H2A.Z are less rigid than the nucleosomes containing H2A (Lewis et al., 2021), which is probably due to the different tail of H2A.Z between plants and animals (Supplementary Fig. S15). The intermediate rigidity of plant nucleosomes containing H2A.Z is consistent with the distribution of H2A.Z in both euchromatin and facultative heterochromatin, supporting its divergent effects on gene transcription in plants (Gomez-Zambrano et al., 2019; Jamge et al., 2023; Wu et al., 2024). Given that H2A.Z is involved in various stress response processes (March-Diaz et al., 2008; Kumar and Wigge, 2010; Smith et al., 2010; Talbert and Henikoff, 2014; Berriri et al., 2016; Cortijo et al., 2017; Sura et al., 2017; Miao et al., 2024), the intermediate state of nucleosomes containing H2A.Z is probably conductive for plants to adapt to environmental changes. Future investigation of the replacement between H2A.Z and H2A or other variants under stress is important to achieve a thorough understanding of the role of H2A.Z in plants.

## Methods

### Plant materials and growth conditions

The *h2a* quadruple mutant and *h2a.w* triple mutant were generated using the CRISPR/Cas9 system in *Arabidopsis thaliana* Columbia ecotype by Biorun Biosciences (Wuhan, China). The *h2a.z* triple mutant was previously reported (Sun et al., 2022). Primers used for identification are listed in Supplementary Table S1. Seeds sterilized with ethyl alcohol were sowed on agar-solidified Murashige & Skoog (MS) medium (M0255; Duchefa, Haarlem, the Netherlands) supplemented with 0.9% (w/v) sucrose. Plants were grown in a reach-in growth chamber at 22 ℃ under either a long-day (16 h light/8 h dark) or short-day (8 h light/16 h dark) photoperiods.

### RNA-seq analysis

Total RNA was extracted from 200 mg 14-d-old seedlings grown under LD conditions using the RNAprep pure Plant Kit (DP432; Tiangen Biotech, Beijing, China). Sequencing libraries were prepared using the KAPA stranded mRNA-Seq Kit for the Illumina^®^ Platform (KR0960-v.5.17; Kapa Biosystems, Wilmington, MA, USA), and sequencing were performed on an Illumina Novaseq 6000 instrument via NEO Bio (Shanghai, China).

Raw reads were processed using Cutadapt v4.4 (Martin, 2011) to remove low-quality bases and sequencing adapters. The trimmed reads were mapped to the reference genomes (TAIR10) using HISAT2 v2.2.1 (Kim et al., 2015), with chloroplast and mitochondrial reads excluded. SAMtools v1.7 software was used to remove low-quality reads (*q* < 20) (Li et al., 2009). High-quality reads were normalized and converted to BigWig format using the bamCoverage tool in deepTools v 3.5.3 software (Ramirez et al., 2016). Genomic track files were visualized using the genome browser IGV v2.11.1 (Thorvaldsdottir et al., 2013). Read counts for gene exons were obtained from bam files using featureCounts v2.0.1 (Liao et al., 2014). Differential expression analysis was conducted with the R package DESeq2 v1.36.0 (Love et al., 2014), using a threshold of log2 (Fold change) | ≥ log2 (2) and *P*-*adj* ≤ 0.01 to identify differentially expressed genes (Supplementary Data Set 1). The R toolkit clusterProfiler v4.4.4 was used to perform GO enrichment analysis (Yu et al., 2012). The screening threshold was set to a *p*-value of ≤ 0.01, and the top 20 enriched biological processes were selected for visualization. Transcriptome-related heatmaps were generated using R package ComplexHeatmap v2.12.1 (Gu et al., 2016). Each RNA-seq sample included at least three independent biological replicates.

### ChIP assay

2 g 14-d-old seedlings were harvested and fixed in the buffer containing 10 mM Tris-HCl, 0.4 M sucrose, 1mM EDTA, 1% v/v formaldehyde, and 1 mM PMSF, pH 8.0 at room temperature for 15 minutes. The fixed seedlings were ground in liquid nitrogen, and the nuclei were isolated in the buffer containing 15 mM PIPES, 0.25 M sucrose, 5 mM MgCl_2_, 60 mM KCl, 15 mM NaCl, 1 mM CaCl_2_, 0.9% v/v Triton X-100, and Protease Inhibitor cocktail (Roche) at pH 6.8. The nuclei were resuspended in the buffer containing 15 mM PIPES, 0.25 M sucrose, 5 mM MgCl_2_, 60 mM KCl, 15 mM NaCl, 1 mM CaCl_2_, 0.9% v/v Triton X-100, 1% w/v SDS, and Protease Inhibitor cocktail (Roche) at pH 6.8. Sonication was performed using a Bioruptor™ Plus sonicator (Diagenode, Liège, Belgium) to fragment the chromatin to sizes below 500 bp. Immunoprecipitation was performed using anti-H2A (A3692; ABclonal, Wuhan, China), anti-H2A.Z (Wu et al., 2023), anti-H2A.W, and anti-H3 (ab1791; Abcam, Cambridge, UK) antibodies. The anti-H2A.W antibody was generated in rabbits using the peptide GGRKPPGAPKTKSVS-C as an antigen (Abmart, Shanghai, China). The immunoprecipitation complexes were gradually washed with the following buffers: a low-salt buffer (20 mM Tris-HCl, 150 mM NaCl, 2 mM EDTA, 0.5% v/v Triton X-100, and 0.2% w/v SDS, pH 8.0), a high-salt buffer (20 mM Tris-HCl, 500 mM NaCl, 2 mM EDTA, 0.5% v/v Triton X-100, and 0.2% w/v SDS, pH 8.0), a LiCl buffer (10 mM Tris-HCl, 0.25 M LiCl, 1 mM EDTA, 1% v/v Nonidet P-40, and 1% v/v sodium deoxycholate, pH 8.0), and a Tris-ETDA buffer (10 mM Tris-HCl and 1 mM EDTA, pH 8.0). The immunoprecipitation complexes were eluted twice from the beads using elution buffer (1% w/v SDS and 0.1 M NaHCO_3_) and then incubated overnight at 65 ℃ for reverse cross-linking. Samples were digested with RNase A (2158; TaKaRa, Dalian, China) and proteinase K (25530015, Invitrogen^TM^, Carlsbad, CA, USA). Finally, DNA fragments were extracted using the phenol-choroform-isopentanol method (Liu et al., 2024).

### ChIP-seq analysis

For the sequencing of H2A, H2A.Z, and H2A.W, libraries were prepared using the VAHTS Universal DNA Library Prep Kit (ND607; Vazyme, Nanjing, China). Sequencing was performed on an Illumina NovaSeq 6000 platform by NEO-BIO (Shanghai, China). For sequencing data analysis, reads trimmed by Cutadapt (Martin, 2011) were aligned to the Arabidopsis reference genome (TAIR10) using BOWTIE2 v.2.5.1 (Langmead and Salzberg, 2012). Low mapping quality reads (*q*<20) and potential PCR duplicates were removed using SAMtools v.1.17 (Li et al., 2009). The enrichment levels (Reads Per Kilobase per Million mapped reads, RPKM) of H3K36me3, H3K9me2, H2A, H2A.Z, and H2A.W were calculated using bamCoverage tool of deepTools v.3.5.3 (Ramirez et al., 2016). The corresponding enrichment peaks were obtained by comparing the differences between IP and Input groups using MACS2 v2.2.7.1 software (Zhang et al., 2008), with a threshold set at *q* < 0.05 (broad-cutoff). BEDtools v2.31.0 (Quinlan and Hall, 2010) was used to identify intersections among the ChIP peaks of H2A and its variants in both arm and centromere regions. Heatmaps from -1.5Kb to 1.5Kb around peak centers were plotted using computeMatrix and plotHeatmap tools in deepTools v3.5.3 (Ramirez et al., 2016).

H3K36me3, H3K9me2, and publicly available H2A.W (Yelagandula et al., 2014) BigWig files were used to distinguish chromosome arm regions from centromeric regions (Supplementary Data Set 3). We identified potential replacement sites for all six possible substitution scenarios (H2A replacing H2A.Z, H2A replacing H2A.W, H2A.Z replacing H2A, H2A.Z replacing H2A.W, H2A.W replacing H2A, and H2A.W replacing H2A.Z). For example, in the case of H2A.Z being replaced by H2A, the mutant *h2a.z*, which lacks the histone variant H2A.Z, serves as a model. In this mutant, sites that are typically enriched with H2A.Z but not with H2A in wild-type plants are the most likely candidates for substitution. These sites were subsequently analyzed to determine the distribution of H2A and H2A.W.

### Preparation of nucleosomes containing Arabidopsis histones and native DNA

All the 147 bp DNA fragments (601, 17.1, 17.2, 15.1.1, 15.1.2, 15.2.1, 15.2.2) were amplified from synthetic DNA templates (Table 1) by using the primers listed in Supplementary Table S1. Arabidopsis histone dimers (H2A-H2B, H2A.Z-H2B, and H2A.W-H2B) and the H3.1-H4 tetramer were co-expressed in *Escherichia coli* Rosetta2 (DE3) cells. The purification of histone dimers and tetramers, the assembly of histone octamers, and nucleosome reconstitution followed previously established protocols (Du et al., 2020). Reconstituted nucleosomes were analyzed using a 5% native polyacrylamide gel in 0.5×TBE buffer (50 mM Tris, 45 mM boric acid, and 1 mM EDTA).

### Cryo-EM grid preparation and data collection

Quantifoil R 1.2/1.3 300 mesh Au grids were glow-discharged in a Pelco easiGlow™ (Ted Pella) at 0.39 mbar, and 15 mA for 60 s. Cross-linked nucleosomes were prepared by incubating the sample with 0.05%-0.1% glutaraldehyde on ice for 15–20 minutes. The nucleosome sample, at a concentration of ∼1 mg/mL, was prepared in 10 mM HEPES-Na (pH 7.4) with 1 mM DTT. A volume of 3.5 μl sample was added to grid and frozen on a Thermo Fisher Scientific™ Vitrobot mark IV, with the relative humidity at 95% at 6 °C. The blot force was set to 1 and the blot time to 3 s, with liquid ethane used as the cryogen.

Grids were loaded onto the Thermo Fisher Scientific™ Titan Krios equipped with a Gatan K3 camera, operated at 300 keV. Data collection was performed using Thermo Scientific™ Smart EPU Software in the super-resolution mode, with a defocus range set between −0.9 and −1.8 μm. The total electron dose of each micrograph stack was 50 e−/Å^2^, dose-fractionated into 40 frames.

### Cryo-EM data processing

The movie stacks of nucleosomes were subjected to dose-weighted motion correction by MotionCor2 (Zheng et al., 2017) and Contrast Transfer Function estimation by CTFFIND4 (Rohou and Grigorieff, 2015). Reference-free 2D classification, initial model Reconstruction, and 3D classification were performed using RELION-v3 (Zivanov et al., 2018). Particles selected through iterative 2D and 3D classifications in RELION were subsequently transferred to cryoSPARC-v4 (Punjani et al., 2017) for further refinement. After 2D classification, Hetero Refinement and Homo Refinement in cryoSPARC, the refined map based on C1 symmetry was respectively reconstructed to 3.34 Å for 147,977 particles of 152-H2A-NCP, 2.93 Å for 192,738 particles of 152-H2A.Z-NCP, 3.05 Å for 119,108 particles of 152-H2A.W-NCP, 3.44 Å for 119,108 particles of 601-H2A-NCP, 3.12 Å for 114,381 particles of 601-H2A.Z-NCP. The 3D reconstruction map imposed C2 symmetry was at a resolution of 3.15 Å for 152-H2A-NCP, 2.75 Å for 152-H2A.Z-NCP, 2.81 Å for 152-H2A.W-NCP, 3.14 Å for 601-H2A-NCP, 2.87 Å for 601-H2A.Z-NCP, respectively. The local resolution estimates were calculated by cryoSPARC.

### Model building, refinement, and validation

Atom models of plant nucleosomes in this study were built using the animal H2A nucleosome structure (PDB ID: 1AOI) as a reference. Briefly, the reference model was fitted into the density maps of each nucleosome using UCSF Chimera (Pettersen et al., 2004). Subsequent real-space refinement was carried out in Phenix (Liebschner et al., 2019). Details building of each model were conducted in Coot (Emsley et al., 2010), and validation was conducted in Phenix. Visualization and rendering of all nucleosome maps were prepared using UCSF ChimeraX (Goddard et al., 2018), while the atom models were prepared using PyMOL-open-source (Delano, 2002). Cryo-EM data collection, 3D reconstruction, model refinement, and validation statistics are summarized in Table 2.

## Data availability

All data supporting the findings of this study are included in this manuscript and supplementary materials. The RNA-seq data and ChIP-seq data are available online (https://www.ncbi.nlm.nih.gov/geo/; accession numbers: GSE283046 and GSE283047). The atomic coordinates and electron microscopy density maps for these structures have been deposited in the PDB (http://www.rcsb.org) and the Electron Microscopy Data Bank (EMDB; https://www.ebi.ac.uk/pdbe/emdb/). The accession numbers are as follows: 152-H2A-nucleosome (C1 Map: EMD-62035, C2 Map: EMD-62036, PDB: 9K40), 152-H2A.Z-nucleosome (C1 Map: EMD-62033, C2 Map: EMD-62034, PDB: 9K3Z), 152-H2A.W-nucleosome (C1 Map: EMD-62037, C2 Map: EMD-62038, PDB: 9K41), 601-H2A-nucleosome (C1 Map: EMD-62039, C2 Map: EMD-62040, PDB: 9K42), 601-H2A.Z-nucleosome (C1 Map: EMD-62041, C2 Map: EMD-62042, PDB: 9K43).

## Author Contributions

A.D. and J.L. conceived the study. A.D. and B.L. supervised the plant experiments. J.Z., J.L. and G.L. supervised the Cryo-EM experiments. Y.W. performed protein expression, purification, structure determination, modeling, and data analysis. J.Wu, S.Y., Q.L and X.Z. performed the plant materials identifications, gene expression, western blotting analysis, and library construction for sequencing. X.L. and J.Wang analyzed the sequencing data. B.L., J.Wu, W.-H. S. and A.D. wrote the manuscript. A.D. edited the manuscript. All authors read, revised, and approved the manuscript.

## Acknowledgments

We thank Dr. Yuda Fang for providing the *h2a.z* triple mutant. This work was supported by the National Key Research and Development Program of China (2024YFE0105100) and the National Natural Science Foundation of China (NSFC31930017, NSFC31800207 and NSFC32100453). We thank the staff members of the Cryo-EM System at the National Facility for Protein Science in Shanghai (NFPS), Shanghai Advanced Research Institute, Chinese Academy of Sciences, and State Key Laboratory of Genetic Engineering, School of Life Sciences, Fudan University for providing technical support and assistance in data collection.

## References

Armeev, G.A., Kniazeva, A.S., Komarova, G.A., Kirpichnikov, M.P., and Shaytan, A.K. (2021). Histone dynamics mediate DNA unwrapping and sliding in nucleosomes. Nat Commun 12, 2387.

Benoit, M., Simon, L., Desset, S., Duc, C., Cotterell, S., Poulet, A., Le Goff, S., Tatout, C., and Probst, A.V. (2019). Replication-coupled histone H3.1 deposition determines nucleosome composition and heterochromatin dynamics during Arabidopsis seedling development. New Phytol 221, 385–398.

Berriri, S., Gangappa, S.N., and Kumar, S.V. (2016). SWR1 Chromatin-Remodeling Complex Subunits and H2A.Z Have Non-overlapping Functions in Immunity and Gene Regulation in Arabidopsis. Mol Plant 9, 1051–1065.

Bourguet, P., Picard, C.L., Yelagandula, R., Pelissier, T., Lorkovic, Z.J., Feng, S., Pouch-Pelissier, M.N., Schmucker, A., Jacobsen, S.E., Berger, F., and Mathieu, O. (2021). The histone variant H2A.W and linker histone H1 co-regulate heterochromatin accessibility and DNA methylation. Nat Commun 12, 2683.

Cortijo, S., Charoensawan, V., Brestovitsky, A., Buning, R., Ravarani, C., Rhodes, D., van Noort, J., Jaeger, K.E., and Wigge, P.A. (2017). Transcriptional Regulation of the Ambient Temperature Response by H2A.Z Nucleosomes and HSF1 Transcription Factors in Arabidopsis. Mol Plant 10, 1258–1273.

Delano, W.L. (2002). PyMOL: An Open-Source Molecular Graphics Tool. CP4 Newsl, Protein Crystallogr.

Du, K., Luo, Q., Yin, L., Wu, J., Liu, Y., Gan, J., Dong, A., and Shen, W.H. (2020). OsChz1 acts as a histone chaperone in modulating chromatin organization and genome function in rice. Nat Commun 11, 5717.

Emsley, P., Lohkamp, B., Scott, W.G., and Cowtan, K. (2010). Features and development of Coot. Acta Crystallogr D Biol Crystallogr 66, 486–501.

Foroozani, M., Holder, D.H., and Deal, R.B. (2022). Histone Variants in the Specialization of Plant Chromatin. Annu Rev Plant Biol 73, 149–172.

Goddard, T.D., Huang, C.C., Meng, E.C., Pettersen, E.F., Couch, G.S., Morris, J.H., and Ferrin, T.E. (2018). UCSF ChimeraX: Meeting modern challenges in visualization and analysis. Protein Sci 27, 14–25.

Gomez-Zambrano, A., Merini, W., and Calonje, M. (2019). The repressive role of Arabidopsis H2A.Z in transcriptional regulation depends on AtBMI1 activity. Nat Commun 10, 2828.

Gu, Z., Eils, R., and Schlesner, M. (2016). Complex heatmaps reveal patterns and correlations in multidimensional genomic data. Bioinformatics 32, 2847–2849.

Hirai, S., Tomimatsu, K., Miyawaki-Kuwakado, A., Takizawa, Y., Komatsu, T., Tachibana, T., Fukushima, Y., Takeda, Y., Negishi, L., Kujirai, T., Koyama, M., Ohkawa, Y., and Kurumizaka, H. (2022). Unusual nucleosome formation and transcriptome influence by the histone H3mm18 variant. Nucleic Acids Res 50, 72–91.

Iouzalen, N., Moreau, J., and Mechali, M. (1996). H2A.ZI, a new variant histone expressed during Xenopus early development exhibits several distinct features from the core histone H2A. Nucleic Acids Res 24, 3947–3952.

Jamge, B., Lorkovic, Z.J., Axelsson, E., Osakabe, A., Shukla, V., Yelagandula, R., Akimcheva, S., Kuehn, A.L., and Berger, F. (2023). Histone variants shape chromatin states in Arabidopsis. Elife 12.

Kawashima, T., Lorkovic, Z.J., Nishihama, R., Ishizaki, K., Axelsson, E., Yelagandula, R., Kohchi, T., and Berger, F. (2015). Diversification of histone H2A variants during plant evolution. Trends Plant Sci 20, 419–425.

Kim, D., Langmead, B., and Salzberg, S.L. (2015). HISAT: a fast spliced aligner with low memory requirements. Nat Methods 12, 357–360.

Kumar, S.V., and Wigge, P.A. (2010). H2A.Z-containing nucleosomes mediate the thermosensory response in Arabidopsis. Cell 140, 136–147.

Langmead, B., and Salzberg, S.L. (2012). Fast gapped-read alignment with Bowtie 2. Nat Methods 9, 357–359.

Lei, B., Capella, M., Montgomery, S.A., Borg, M., Osakabe, A., Goiser, M., Muhammad, A., Braun, S., and Berger, F. (2021). A Synthetic Approach to Reconstruct the Evolutionary and Functional Innovations of the Plant Histone Variant H2A.W. Curr Biol 31, 182–191 e185.

Lewis, T.S., Sokolova, V., Jung, H., Ng, H., and Tan, D.Y. (2021). Structural basis of chromatin regulation by histone variant H2A.Z. Nucleic Acids Research 49, 11379–11391.

Li, H., Handsaker, B., Wysoker, A., Fennell, T., Ruan, J., Homer, N., Marth, G., Abecasis, G., Durbin, R., and Genome Project Data Processing, S. (2009). The Sequence Alignment/Map format and SAMtools. Bioinformatics 25, 2078–2079.

Liao, Y., Smyth, G.K., and Shi, W. (2014). featureCounts: an efficient general purpose program for assigning sequence reads to genomic features. Bioinformatics 30, 923–930.

Liebschner, D., Afonine, P.V., Baker, M.L., Bunkoczi, G., Chen, V.B., Croll, T.I., Hintze, B., Hung, L.W., Jain, S., McCoy, A.J., Moriarty, N.W., Oeffner, R.D., Poon, B.K., Prisant, M.G., Read, R.J., Richardson, J.S., Richardson, D.C., Sammito, M.D., Sobolev, O.V., Stockwell, D.H., Terwilliger, T.C., Urzhumtsev, A.G., Videau, L.L., Williams, C.J., and Adams, P.D. (2019). Macromolecular structure determination using X-rays, neutrons and electrons: recent developments in Phenix. Acta Crystallogr D Struct Biol 75, 861–877.

Liu, B., Li, C.Z., Li, X., Wang, J.C., Xie, W.H., Woods, D.P., Li, W.Y., Zhu, X.Y., Yang, S.M., Dong, A.W., and Amasino, R.M. (2024). The H3K4 demethylase JMJ1 is required for proper timing of flowering in Brachypodium distachyon. Plant Cell 36, 2729–2745.

Liu, C., Lu, F., Cui, X., and Cao, X. (2010). Histone methylation in higher plants. Annu Rev Plant Biol 61, 395–420.

Lorkovic, Z.J., Park, C., Goiser, M., Jiang, D., Kurzbauer, M.T., Schlogelhofer, P., and Berger, F. (2017). Compartmentalization of DNA Damage Response between Heterochromatin and Euchromatin Is Mediated by Distinct H2A Histone Variants. Curr Biol 27, 1192–1199.

Love, M.I., Huber, W., and Anders, S. (2014). Moderated estimation of fold change and dispersion for RNA-seq data with DESeq2. Genome Biol 15, 550.

Lowary, P.T., and Widom, J. (1998). New DNA sequence rules for high affinity binding to histone octamer and sequence-directed nucleosome positioning. J Mol Biol 276, 19–42.

Luger, K., Mader, A.W., Richmond, R.K., Sargent, D.F., and Richmond, T.J. (1997). Crystal structure of the nucleosome core particle at 2.8 A resolution. Nature 389, 251–260.

Malik, H.S., and Henikoff, S. (2003). Phylogenomics of the nucleosome. Nat Struct Biol 10, 882–891.

March-Diaz, R., Garcia-Dominguez, M., Lozano-Juste, J., Leon, J., Florencio, F.J., and Reyes, J.C. (2008). Histone H2A.Z and homologues of components of the SWR1 complex are required to control immunity in Arabidopsis. Plant J 53, 475–487.

Martin, M. (2011). Cutadapt removes adapter sequences from high-throughput sequencing reads. EMBnet.journal. 17, 10.

Marzluff, W.F., Wagner, E.J., and Duronio, R.J. (2008). Metabolism and regulation of canonical histone mRNAs: life without a poly(A) tail. Nat Rev Genet 9, 843–854.

Miao, R., Zhang, Y., Liu, X., Yuan, Y., Zang, W., Li, Z., Yan, X., Pang, Q., and Zhang, A. (2024). Histone variant H2A.Z is required for plant salt response by regulating gene transcription. Plant Cell Environ 47, 2693–2709.

Osakabe, A., Takizawa, Y., Horikoshi, N., Hatazawa, S., Negishi, L., Sato, S., Berger, F., Kakutani, T., and Kurumizaka, H. (2024). Molecular and structural basis of the chromatin remodeling activity by Arabidopsis DDM1. Nat Commun 15, 5187.

Osakabe, A., Jamge, B., Axelsson, E., Montgomery, S.A., Akimcheva, S., Kuehn, A.L., Pisupati, R., Lorkovic, Z.J., Yelagandula, R., Kakutani, T., and Berger, F. (2021). The chromatin remodeler DDM1 prevents transposon mobility through deposition of histone variant H2A.W. Nat Cell Biol 23, 391–400.

Pettersen, E.F., Goddard, T.D., Huang, C.C., Couch, G.S., Greenblatt, D.M., Meng, E.C., and Ferrin, T.E. (2004). UCSF Chimera--a visualization system for exploratory research and analysis. J Comput Chem 25, 1605–1612.

Probst, A.V. (2022). Deposition and eviction of histone variants define functional chromatin states in plants. Curr Opin Plant Biol 69, 102266.

Punjani, A., Rubinstein, J.L., Fleet, D.J., and Brubaker, M.A. (2017). cryoSPARC: algorithms for rapid unsupervised cryo-EM structure determination. Nat Methods 14, 290–296.

Quinlan, A.R., and Hall, I.M. (2010). BEDTools: a flexible suite of utilities for comparing genomic features. Bioinformatics 26, 841–842.

Ramirez, F., Ryan, D.P., Gruning, B., Bhardwaj, V., Kilpert, F., Richter, A.S., Heyne, S., Dundar, F., and Manke, T. (2016). deepTools2: a next generation web server for deep-sequencing data analysis. Nucleic Acids Res 44, W160–165.

Rangasamy, D., Berven, L., Ridgway, P., and Tremethick, D.J. (2003). Pericentric heterochromatin becomes enriched with H2A.Z during early mammalian development. EMBO J 22, 1599–1607.

Rohou, A., and Grigorieff, N. (2015). CTFFIND4: Fast and accurate defocus estimation from electron micrographs. Journal of Structural Biology 192, 216–221.

Rosa, M., Von Harder, M., Cigliano, R.A., Schlogelhofer, P., and Mittelsten Scheid, O. (2013). The Arabidopsis SWR1 chromatin-remodeling complex is important for DNA repair, somatic recombination, and meiosis. Plant Cell 25, 1990–2001.

Roudier, F., Ahmed, I., Berard, C., Sarazin, A., Mary-Huard, T., Cortijo, S., Bouyer, D., Caillieux, E., Duvernois-Berthet, E., Al-Shikhley, L., Giraut, L., Despres, B., Drevensek, S., Barneche, F., Derozier, S., Brunaud, V., Aubourg, S., Schnittger, A., Bowler, C., Martin-Magniette, M.L., Robin, S., Caboche, M., and Colot, V. (2011). Integrative epigenomic mapping defines four main chromatin states in Arabidopsis. EMBO J 30, 1928–1938.

Smith, A.P., Jain, A., Deal, R.B., Nagarajan, V.K., Poling, M.D., Raghothama, K.G., and Meagher, R.B. (2010). Histone H2A.Z regulates the expression of several classes of phosphate starvation response genes but not as a transcriptional activator. Plant Physiol 152, 217–225.

Sun, A., Yin, C., Ma, M., Zhou, Y., Zheng, X., Tu, X., and Fang, Y. (2022). Feedback regulation of auxin signaling through the transcription of H2A.Z and deposition of H2A.Z to SMALL AUXIN UP RNAs in Arabidopsis. New Phytol 236, 1721–1733.

Sura, W., Kabza, M., Karlowski, W.M., Bieluszewski, T., Kus-Slowinska, M., Paweloszek, L., Sadowski, J., and Ziolkowski, P.A. (2017). Dual Role of the Histone Variant H2A.Z in Transcriptional Regulation of Stress-Response Genes. Plant Cell 29, 791–807.

Suto, R.K., Clarkson, M.J., Tremethick, D.J., and Luger, K. (2000). Crystal structure of a nucleosome core particle containing the variant histone H2A.Z. Nat Struct Biol 7, 1121–1124.

Talbert, P.B., and Henikoff, S. (2014). Environmental responses mediated by histone variants. Trends Cell Biol 24, 642–650.

Talbert, P.B., Ahmad, K., Almouzni, G., Ausio, J., Berger, F., Bhalla, P.L., Bonner, W.M., Cande, W.Z., Chadwick, B.P., Chan, S.W., Cross, G.A., Cui, L., Dimitrov, S.I., Doenecke, D., Eirin-Lopez, J.M., Gorovsky, M.A., Hake, S.B., Hamkalo, B.A., Holec, S., Jacobsen, S.E., Kamieniarz, K., Khochbin, S., Ladurner, A.G., Landsman, D., Latham, J.A., Loppin, B., Malik, H.S., Marzluff, W.F., Pehrson, J.R., Postberg, J., Schneider, R., Singh, M.B., Smith, M.M., Thompson, E., Torres-Padilla, M.E., Tremethick, D.J., Turner, B.M., Waterborg, J.H., Wollmann, H., Yelagandula, R., Zhu, B., and Henikoff, S. (2012). A unified phylogeny-based nomenclature for histone variants. Epigenetics Chromatin 5, 7.

Thorvaldsdottir, H., Robinson, J.T., and Mesirov, J.P. (2013). Integrative Genomics Viewer (IGV): high-performance genomics data visualization and exploration. Brief Bioinform 14, 178–192.

Wu, J., Liu, B., and Dong, A. (2024). Interplay between histone variants and chaperones in plants. Curr Opin Plant Biol 80, 102551.

Wu, J., Yang, Y., Wang, J., Wang, Y., Yin, L., An, Z., Du, K., Zhu, Y., Qi, J., Shen, W.H., and Dong, A. (2023). Histone chaperones AtChz1A and AtChz1B are required for H2A.Z deposition and interact with the SWR1 chromatin-remodeling complex in Arabidopsis thaliana. New Phytol 239, 189–207.

Xue, M., Zhang, H., Zhao, F., Zhao, T., Li, H., and Jiang, D. (2021). The INO80 chromatin remodeling complex promotes thermomorphogenesis by connecting H2A.Z eviction and active transcription in Arabidopsis. Mol Plant 14, 1799–1813.

Yelagandula, R., Stroud, H., Holec, S., Zhou, K., Feng, S., Zhong, X., Muthurajan, U.M., Nie, X., Kawashima, T., Groth, M., Luger, K., Jacobsen, S.E., and Berger, F. (2014). The histone variant H2A.W defines heterochromatin and promotes chromatin condensation in Arabidopsis. Cell 158, 98–109.

Yi, H., Sardesai, N., Fujinuma, T., Chan, C.W., Veena, and Gelvin, S.B. (2006). Constitutive expression exposes functional redundancy between the Arabidopsis histone H2A gene HTA1 and other H2A gene family members. Plant Cell 18, 1575–1589.

Yu, G., Wang, L.G., Han, Y., and He, Q.Y. (2012). clusterProfiler: an R package for comparing biological themes among gene clusters. OMICS 16, 284–287.

Zhang, H., Gu, Z., Zeng, Y., and Zhang, Y. (2024). Mechanism of heterochromatin remodeling revealed by the DDM1 bound nucleosome structures. Structure 32, 1222–1230 e1224.

Zhang, Y., Liu, T., Meyer, C.A., Eeckhoute, J., Johnson, D.S., Bernstein, B.E., Nusbaum, C., Myers, R.M., Brown, M., Li, W., and Liu, X.S. (2008). Model-based analysis of ChIP-Seq (MACS). Genome Biol 9, R137.

Zheng, S.Q., Palovcak, E., Armache, J.-P., Verba, K.A., Cheng, Y., and Agard, D.A. (2017). MotionCor2: anisotropic correction of beam-induced motion for improved cryo-electron microscopy. Nature Methods 14, 331–332.

Zivanov, J., Nakane, T., Forsberg, B.O., Kimanius, D., Hagen, W.J., Lindahl, E., and Scheres, S.H. (2018). New tools for automated high-resolution cryo-EM structure determination in RELION-3. Elife 7.

